# Mutation nsp6 L232F associated with MERS-CoV zoonotic transmission confers higher viral replication in human respiratory tract cultures ex-vivo

**DOI:** 10.1101/2023.03.27.534490

**Authors:** Ray TY So, Daniel KW Chu, Kenrie PY Hui, Chris KP Mok, Sumana Sanyal, John M Nicholls, John C. W. Ho, Man-chun Cheung, Ka-chun Ng, Hin-Wo Yeung, Michael CW Chan, Leo LM Poon, Jincun Zhao, Malik Peiris

## Abstract

Middle East respiratory syndrome coronavirus (MERS-CoV) causes zoonotic disease. Dromedary camels are the source of zoonotic infection. We identified a mutation of amino acid leucine to phenylalanine in the codon 232 position of the non-structural protein 6 (nsp6) (nsp6 L232F) that is repeatedly associated with zoonotic transmission. We generated a pair of isogenic recombinant MERS-CoV with nsp6 232L and 232F residues, respectively, and showed that the nsp6 L232F mutation confers higher replication competence in ex-vivo culture of human nasal and bronchial tissues and in lungs of mice experimentally infected in-vivo. Mechanistically, the nsp6 L232F mutation appeared to modulate autophagy and was associated with higher exocytic virus egress, while innate immune responses and zippering activity of the endoplasmic reticulum remained unaffected. Our study suggests that MERS-CoV nsp6 may contribute to viral adaptation to humans. This highlights the importance of continued surveillance of MERS-CoV in both camels and humans.

**Importance:** Viral host adaptation plays an important role in zoonotic transmission of coronaviruses. For MERS-CoV that widely circulates in dromedary camels from Arabian Peninsula, camel-to human transmissions are ongoing, raising the possibility of human adaptive evolution for MERS-CoV. Here, we analysed MERS-CoV sequences and identified an amino acid mutation L232F in nsp6 to occur repeatedly in human MERS-CoV over the years since the first outbreak in 2012. We found the nsp6 L232F confers increase viral replication in-vitro, in ex-vivo upper human respiratory tract cultures and in mice, using a reverse genetics approach. Our results showed the nsp6 L232F may be advantageous for MERS-CoV to replicate in humans. This study highlighted a human adaptation of MERS-CoV and a need for continued surveillance of MERS-CoV to identify any further adaptations in humans, which may be relevant to the pandemic potential of MERS-CoV.

## Introduction

Middle East respiratory syndrome coronavirus (MERS-CoV) is an emerging pathogen that was first recognized in 2012 as a cause of severe acute respiratory diseases in humans [1]. Although another novel coronavirus, severe acute respiratory coronavirus 2 (SARS-CoV-2) emerged in 2019 to cause a pandemic (COVID-19), MERS-CoV remains a potential pandemic threat and remains a research priority [2-4]. As of December 2022, MERS-CoV had led to 2603 confirmed human cases, 935 of which were fatal [5]. Dromedary camels are the source of zoonotic human infection [6, 7]. MERS-CoVs infected dromedary camels are found in the Arabian Peninsula (MERS-CoV clades A and B), in Africa (clade C) and Central Asia (phylogenetics undefined so far). Zoonotic disease has only been reported in the Arabian Peninsula [4, 8]. The exact mode of inter-species transmission remains unknown but entry via the respiratory or gastrointestinal tracts are the plausible transmission routes and may occur via direct or indirect camel contact [8]. There is no evidence of sustained transmission of MERS-CoV in the local community, although nosocomial outbreaks have occurred in Saudi Arabia (2014, 2015, 2016, and 2018) and South Korea (2015), which may sometimes exceed one hundred or more individuals [9-12]. Such human-to-human transmission occurred mainly between patients, healthcare workers and visitors.

Genetic adaptation associated with inter-species transmission of viruses have been documented in severe acute respiratory syndrome coronavirus (SARS-CoV-1) and avian influenza virus [13, 14]. SARS-CoV-2 also demonstrated genetic adaptation (e.g. D614G amino acid substitution in spike protein) associated with increased transmissibility and fitness within humans [15]. Host adaptation mutations of MERS-CoV in humans remains poorly understood. Deletions in genes encoding accessory proteins have been reported, including one from a human MERS-CoV outbreak cluster and another one from an individual patient, but such variants occurred only on a single occasion and eventually died out in human transmission [16, 17]. It is important to investigate host adaptive mutations association with zoonotic transmission of MERS-CoV.

In this study, we aimed to identify mutations that were biased to occur more frequently in human rather than in dromedary camel MERS-CoV sequences. We identified a nsp6 L232F mutation that preferentially occurred in human MERS-CoV sequences. Using reverse genetics, we demonstrated that the mutation conferred higher replication competence in the human respiratory tract. MERS-CoV nsp6 was previously shown to interact with different cellular processes, in particular, autophagy restriction and innate immune antagonism [18-20]. Nsp6 has also recently been shown to play an important role in the biogenesis of viral induced double membrane vesicles, known to be a replication organelle for viral RNA synthesis [21]. In attempting to identify the molecular mechanism of the nsp6 L232F adaptive mutation, here we show that the mutation could enhance autophagy during infection. Our study highlighted the need for a continued surveillance of MERS-CoV evolution.

## Results

### Repeated independent emergence of nsp6 L232F mutation in human MERS-CoV from multiple zoonotic introductions suggests convergent evolution

We retrieved a total of 502 MERS-CoV complete and partial (>20kb) genomes from Virus Variation, National Centre for Biotechnology Information (NCBI) database to identify mutations that were preferentially biased to occur in human MERS-CoV. The dataset includes all available MERS-CoV sequences including clade A and B viruses from the Arabian Peninsula and clade C viruses from Africa. Using Fisher’s exact test adjusted with Bonferroni correction, 11 mutations were found to occur significantly more frequently in MERS-CoV sequences from humans than from camels (Table 1). These mutations were found in nsp3, nsp5, nsp6, nsp13, spike and nucleocapsid viral proteins. The mutations with the greatest difference in occurrence between human and camel origin MERS-CoV sequences was the nsp6 L232F mutation, which occurred in 60/236 (25%) of the human sequences but in just 1/266 (0.004%) of the camel MERS-CoV sequences, the camel virus with nsp6 L232F being found in a clade B virus. We generated a phylogenetic tree to visualize the occurrence of the nsp6 mutation in the MERS-CoV phylogeny (Fig. 1, enlarged version in Supplementary Fig.1). The first human MERS-CoV sequence that showed the nsp6 L232F mutation was detected in Saudi Arabia in May 2013, belonging to lineage 2, clade B of the tree. The same nsp6 L232F mutation was also detected to occur independently in multiple human clade B MERS-CoV sequences from lineages 3, 4 and 5. The pattern reveals a convergent human adaptive mutation in MERS-CoV.

**Table 1.**
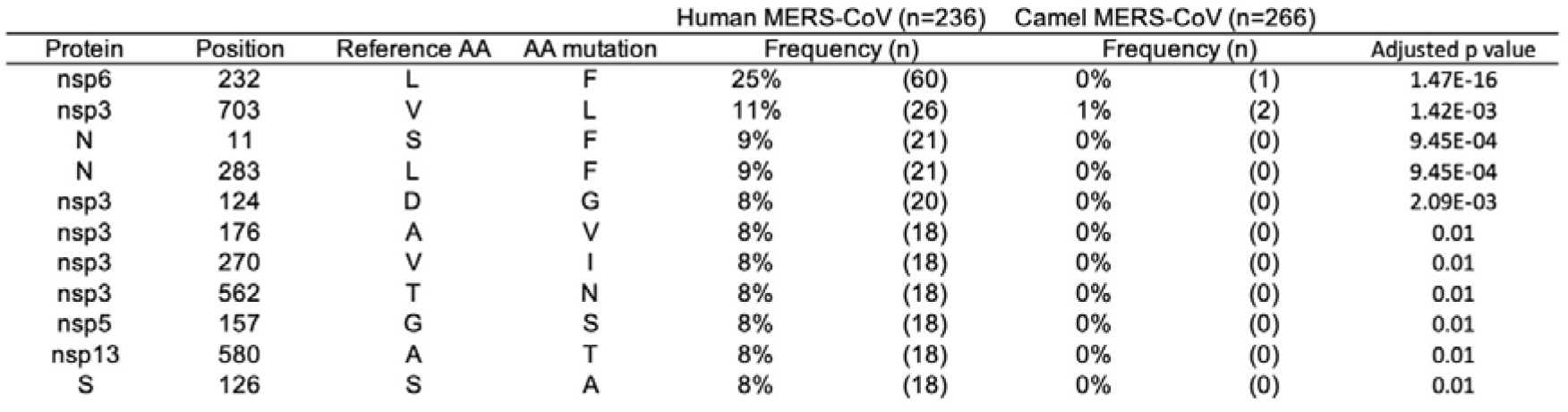
List of amino acid mutations that preferentially occurred in MERS-CoV sequences from human but not camels. The table was generated from a dataset of a total of 502 non-identical MERS-CoV sequences from camels and humans. Each sequence contains 9836 codon position. A cut-off was set to only include amino acid mutation with a frequency of >5% in human sequences and <1% in camel sequences. Test for independence was done by Fisher’s exact test, with the p-value adjusted for multiple testing using conservative Bonferroni correction.

| Protein | Position | Reference AA | AA mutation | Human MERS-CoV (n=236) |  | Camel MERS-CoV (n=266) |  | Adjusted p value |
| --- | --- | --- | --- | --- | --- | --- | --- | --- |
|  |  |  |  | Frequency (%) | Frequency (n) | Frequency (%) | Frequency (n) |  |
| nsp6 | 232 | L | F | 25% | (60) | 0% | (1) | 1.47E-16 |
| nsp3 | 703 | V | L | 11% | (26) | 1% | (2) | 1.42E-03 |
| N | 11 | S | F | 9% | (21) | 0% | (0) | 9.45E-04 |
| N | 283 | L | F | 9% | (21) | 0% | (0) | 9.45E-04 |
| nsp3 | 124 | D | G | 8% | (20) | 0% | (0) | 2.09E-03 |
| nsp3 | 176 | A | V | 8% | (18) | 0% | (0) | 0.01 |
| nsp3 | 270 | V | I | 8% | (18) | 0% | (0) | 0.01 |
| nsp3 | 562 | T | N | 8% | (18) | 0% | (0) | 0.01 |
| nsp5 | 157 | G | S | 8% | (18) | 0% | (0) | 0.01 |
| nsp13 | 580 | A | T | 8% | (18) | 0% | (0) | 0.01 |
| S | 126 | S | A | 8% | (18) | 0% | (0) | 0.01 |

**Figure 1.**
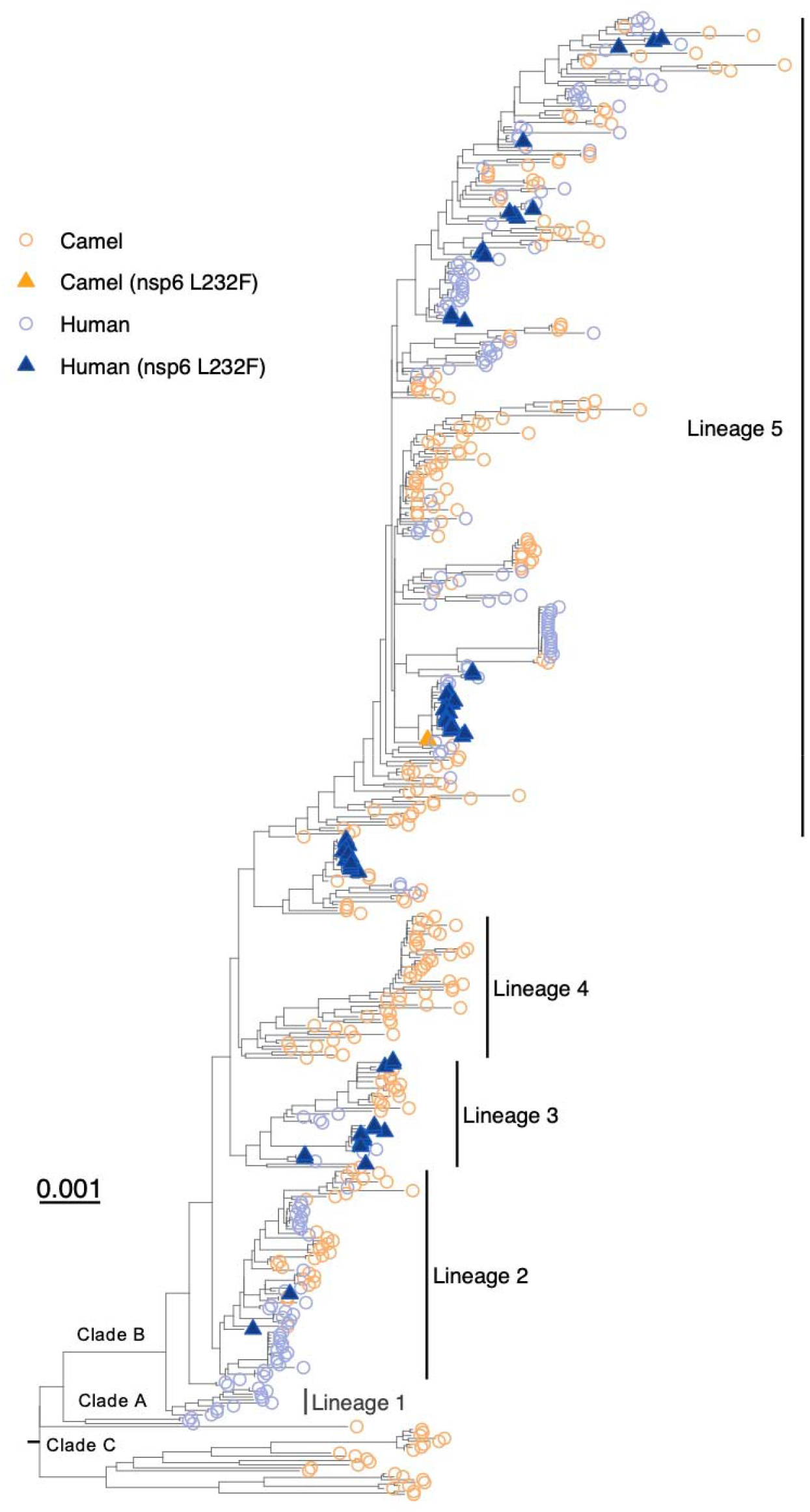
The occurrence of MERS-CoV nsp6 L232F mutation in human and dromedary camel MERS-CoV. A PhyML tree of 502 complete MERS-CoV genomes labelled with the host and the presence of the mutation. Tree was built with IQ-Tree v1.6.12. Yellow color represents camel sequences; blue color represents human sequences. Taxon with a triangle shape indicates the presence of the nsp6 L232F mutation. Scale bar, 0.001 substitution per site.

To ascertain whether the nsp L232F mutation pre-exists as a minor variant within the intra-host genetic diversity in camel MERS-CoV, we analysed the Next Gen Sequencing raw read data of 8 camel swab samples collected in Saudi Arabia (2015-2018, 2 samples per year). None of the samples carried the nsp L232F mutation as a minor variant (coverage of > 10,000 reads in the nsp6 region, Supplementary Table 1). This suggests the mutation is likely to arise after infecting human hosts, instead of being transferred from the camel MERS-CoV quasispecies.

### Isogenic virus clones generated by reverse genetics showed that the nsp6 L232F mutation enhances viral replication in Calu-3 cells, Vero cells and in hDPP4 knockin mice

To study the effect of the nsp6 L232F mutation on viral replication, we used reverse genetics to rescue a pair of isogenic recombinant viruses from a wild-type human clade B MERS-CoV strain, ChinaGD01, that contains the nsp6-232Phe mutation (rGD01-WT) and a mutant isogenic ChinaGD01 that was mutated to carry the camel predominant Leu residue in the nsp6 232 position (rGD01-nsp6 F232L). In Calu-3 cells, both rGD01-WT and rGD01-nsp6 232L viruses replicated similarly in multicycle infections (MOI=0.01) from 24 – 48 hours but rGD01-WT had significantly higher titres at 72 hours post-infection (hpi) (Fig. 2A). In Vero cells, that are deficient in type I interferon (IFN-I) production, rGD01-WT replicated to significantly higher viral titers than rGD01-nsp6 232L at 24 and 48 hpi (MOI of 0.01) (Fig. 2B). Plaque morphology in Vero cells showed that rGD01-WT formed plaques of larger sizes than rGD01-nsp6 232L (fold difference: 1 vs 0.71) (Fig. 2C). The pair of viruses were infected at a MOI of 2 to measure early viral kinetics and again rGD01-WT showed higher viral replication in infectious virus assays as well as higher levels of genomic and subgenomic viral RNA synthesis at 12 and 24 hpi (Fig 2D). A direct growth competition assay between the virus-pair was performed in Vero cells with different ratios of rGD01-WT and rGD01-nsp6 232L ranging from 9:1, 1:1 and 1:9, at a MOI of 0.01 (Fig. 2E). After 3 rounds of serial passage, nsp6 232Phe emerged as the dominant genotype at all infection ratios in Vero cells, supporting the contention that rGD01-WT has higher intrinsic fitness and replication competence.

**Figure 2.**
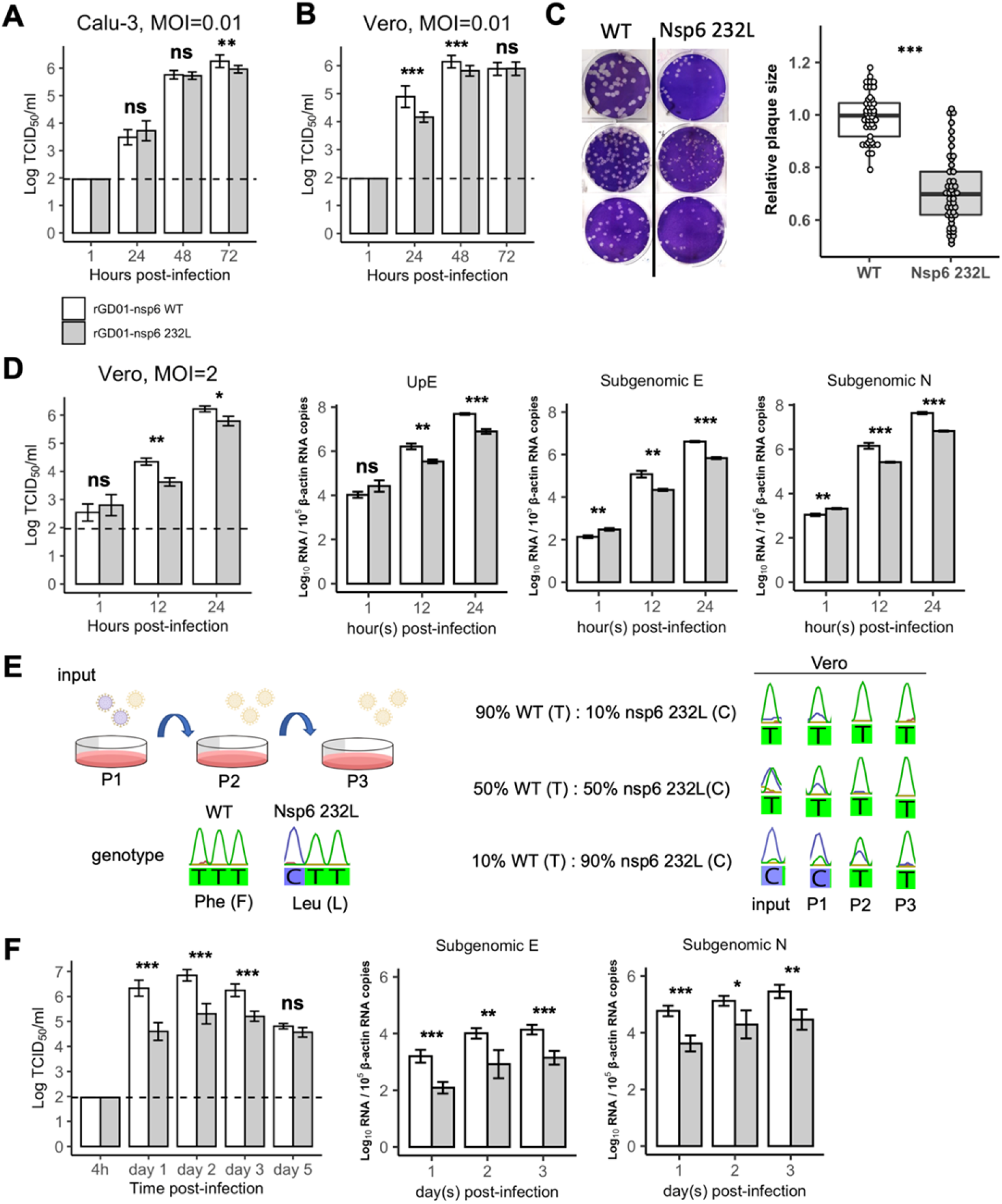
Comparison of virus replication kinetics of rGD01-WT and rGD01-nsp6 232L isogenic MERS-CoV. (A) Calu-3 cell cultures and (B) Vero cell cultures were infected at an MOI of 0.01 and virus titers in culture supernatants were determined by TCID_50_ assay. Assays were performed in triplicates. Data are mean ± s.d. (C) Relative plaque sizes of rGD01-WT and rGD01-nsp6 232L virus were determined in Vero cells. Assays were performed in triplicates. A total of n=46 plaques were analyzed for each virus. The box plot displays median and interquartile range. (D) Early cycles replication kinetics in Vero cells (n=3) at MOI=2 measured for live virus production and RNA synthesis. Assays were performed in triplicates. Data are mean ± s.d. (E) Schematic setup of virus growth competition assay in Vero cells. Vero cells were infected with different ratios (9:1, 1:1 and 1:9) (n=1) of rGD01-WT and rGD01-nsp6 232L virus and serially passaged three times. Genotype of the nsp6 amino acid 232 position was determined by sanger sequencing. (F) Viral titers in the lungs of human DPP4-knockin mice infected intranasally with 1 × 10^4^ pfu of each virus (n=4 for each group). Lung homogenates (n=4) were measured for viral titers by TCID_50_ assay and total RNA extracted to measure subgenomic viral RNA templates by RT-qPCR targeting the envelope (subgenomic E) and nucleocapsid (subgenomic N) gene. Statistical tests were done using two tailed Student’s t test: p ≥0.05 (ns); p <0.05 (*); p <0.01 (**); p <0.001 (***).

Viral kinetics were then compared in the human DPP4 knockin C57BL/6 mouse model that has been previously used for MERS-CoV replication studies [22]. Following intranasal inoculation of 10^4^ plaque forming units (pfu) of each isogenic virus per mouse, lung viral titers were significantly higher in rGD01-WT virus at 1 to 3 days post-infection (dpi), with the greatest difference of 1.71 log_10_ 50% tissue culture infection dose (TCID_50_)/ml observed on day 1 (Fig. 2F). Titer differences between the viruses gradually diminished from day 2 to day 5. Higher subgenomic RNA production was seen in rGD01-WT than in rGD01-nsp6 232L from day 1 to 3 dpi. Overall, these data suggest the nsp6 L232F mutation will confer a higher viral fitness in the mouse respiratory tract.

### Nsp6 L232F mutation enhances viral replication in ex-vivo cultures of the human upper and lower respiratory tract

To evaluate the potential changes of tropism in the human respiratory tract, we assessed the effect of the nsp6 L232F mutation in ex-vivo cultures of nasal, bronchial and lung tissues. We compared the viral replication kinetics of the isogenic viruses at 33°C in nasal turbinate tissues and at 37°C in bronchus and lung tissues. In the nasal tissues, rGD01-WT showed a significant increase in virus titer at 24 hpi, but differences were not significantly different at 48 hpi. Aggregating virus titers at 24-48 hpi using area under the curve (AUC) analysis showed an overall increase in release of infectious virus in nasal tissues (Fig. 3A). Similarly, rGD01-WT showed significantly higher virus titers at 48 to 72 hpi, as well as in the AUC analysis, in ex-vivo cultures of human bronchus (Fig. 3B). No significant differences in virus titers were observed in lung tissues at 24, 48 or 72 hpi (Fig. 3C). Overall, the nsp6 L232F showed enhanced viral replication in human upper respiratory tract and conducting airway tissues.

**Figure 3.**
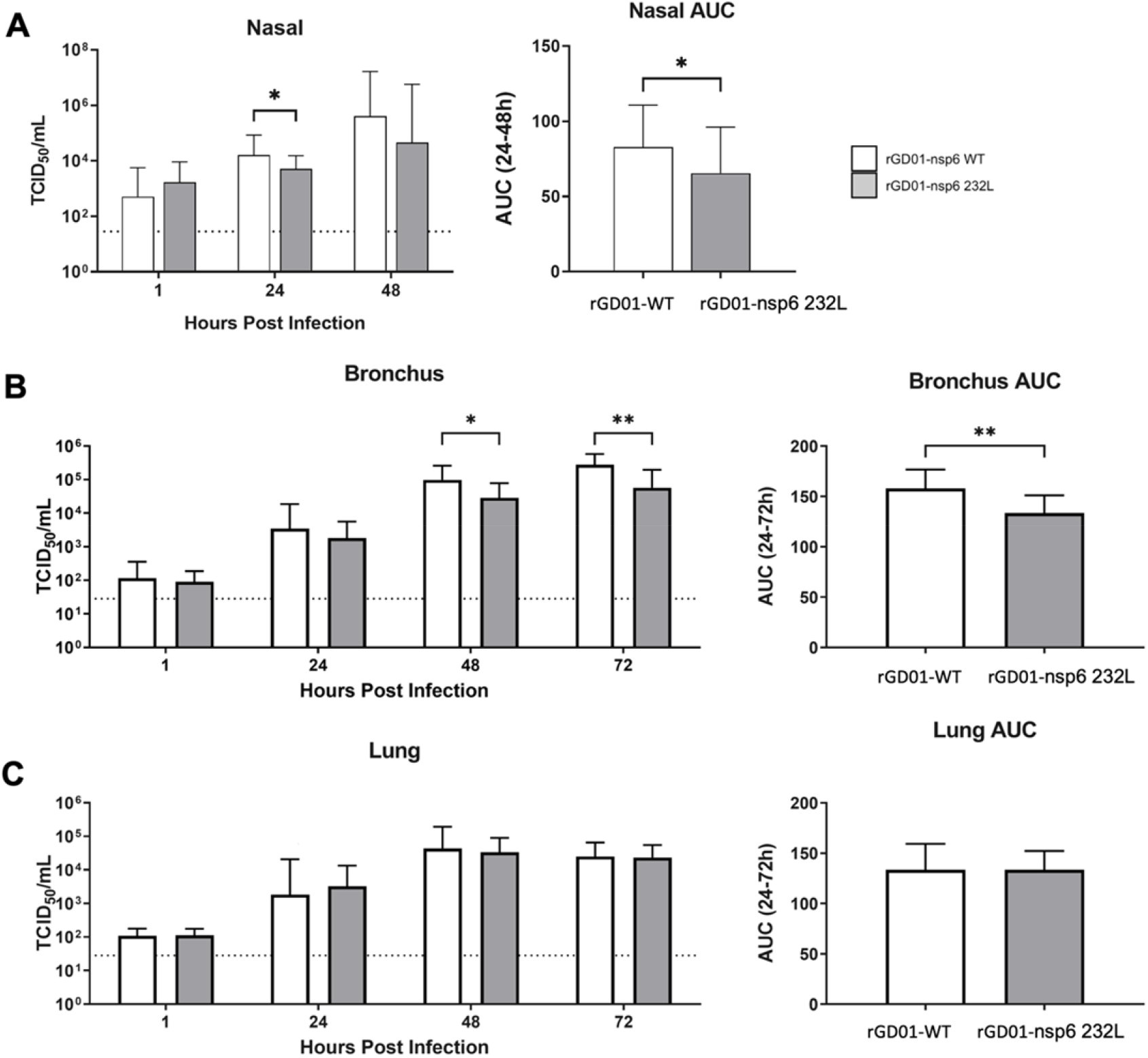
Comparison of virus replication kinetics in ex-vivo cultures of nasal, bronchial and lung tissues. Ex vivo cultures of nasal (A), bronchus (B), and lung (C) tissues were infected with a dose of 10^6^ TCID_50_ of rGD01-WT and rGD01-nsp6 232L viruses. Virus titers in culture supernatants at the indicated timepoints were determined by TCID_50_ assay. Data were generated from three individual tissue donors and represented as mean and standard error. The horizontal dotted line denotes the limit of detection in the TCID_50_ assay. Statistical tests were done using paired t test: P <0.05 (*), P <0.01(**).

### Nsp6 L232F mutation did not affect innate immune antagonism

To explore the molecular basis of how the nsp6 L232F mutation may mediate an enhanced virus replication phenotype, we first tested innate immune responses elicited by infection of Calu-3 cells with the pair of isogenic viruses. Nsp6 has previously been shown to inhibit IFN-I production *in vitro* in an overexpression system [20]. We measured the mRNA expression of a panel of IFN-stimulated genes in Calu-3 cells, at 24 and 48 hpi following infection with each virus at MOI=2. No significant differences in the mRNA expression of IFN-β, TNF-α, IL6, ISG15, RANTES and CXCL-19 (IP-10) were observed at both 24 and 48 hpi (Fig. 4A), viral RNA copies being shown in Fig. 4B. This suggests nsp6 L232F mutation did not alter innate immune antagonism upon infection.

**Figure 4.**
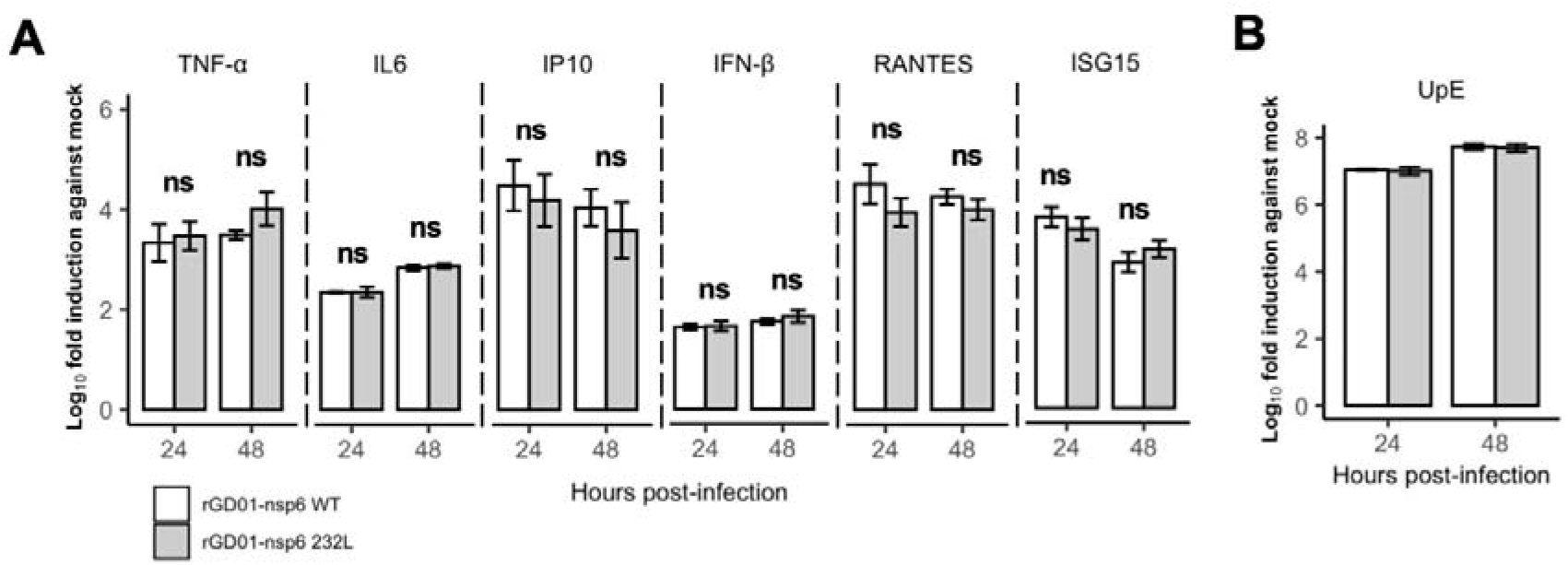
Innate immune gene expression in Calu-3 cells infected with rGD01-WT and rGD01-nsp6 232L viruses (MOI=2). (A) mRNA expression of immune genes (TNF-α, IL6, IP10, IFN-β, RANTES and ISG15) were determined using qPCR from cells infected with rGD01-WT or rGD01-nsp6 232L virus. Mock control was cells mock infected without virus infection. Assays were performed in triplicates. Data are mean ± s.d. (B) Quantification of MERS-CoV RNA copies using RT-qPCR targeting the upstream region of E gene (UpE). Assays were performed in triplicates. Data are mean ± s.d. Statistical tests were done using Student’s t test.

### Nsp6 L232F mutation modulates autophagy

MERS-CoV infection has been shown to restrict autophagic flux contributed by nsp6 [23]. We therefore studied the effect of the nsp6 L232F mutation on the induction and flux of autophagy, as it plays important roles in restricting infection through degradation of viral particles. We infected Vero cells transfected with a tandem fluorescent-tagged LC3 (mRFP-EGFP-LC3) with the isogenic viruses at MOI=0.01 and measured the autophagic compartments (red puncta = autolysosomes (AL); yellow puncta = autophagosomes (AP)) at 24 hpi (Fig. 5A). Infection of both viruses reduced AL puncta, when compared to mock control, suggesting restriction of autophagy from both viruses (Fig. 5B). A moderate reduction of AP puncta was observed in rGD01-WT compared to rGD01-nsp6 232L, indicating less autophagic restriction of the rGD01-WT virus (Fig. 5B). In a parallel approach, we measured total LC3 levels by immunoblotting, where LC3-I (16-18 kDa) corresponds to its cytoplasmic form, and LC3-II (14-16 kDa) corresponds to AP associated LC3. With Bafilomycin A1 (BafA1) treatment which blocks the fusion of AP to AL, rGD01-WT showed an increase in LC3-II/ LC3-I ratio, but not in rGD01-nsp6 232L at 24 hpi (Fig. 5C). This indicated that rGD01-WT infection resulted in reduced autophagic restriction. To visualize the autophagic compartments, we performed electron microscopy on infected Vero cells and showed a higher numbers of multivesicular bodies with rGD01-WT compared to rGD01-nsp6 232L (Fig. 5D). Virus particles were observed in these vesicles, suggesting that an autophagy-derived exocytic pathway may be exploited for cellular egress of MERS-CoV. We observed a larger vesicle size containing higher numbers of virus particles in rGD01-WT compared to rGD01-nsp6 232L infected cells. Overall, we observed a modulation of autophagy and a potential effect on exocytosis for virus egress.

**Figure 5.**
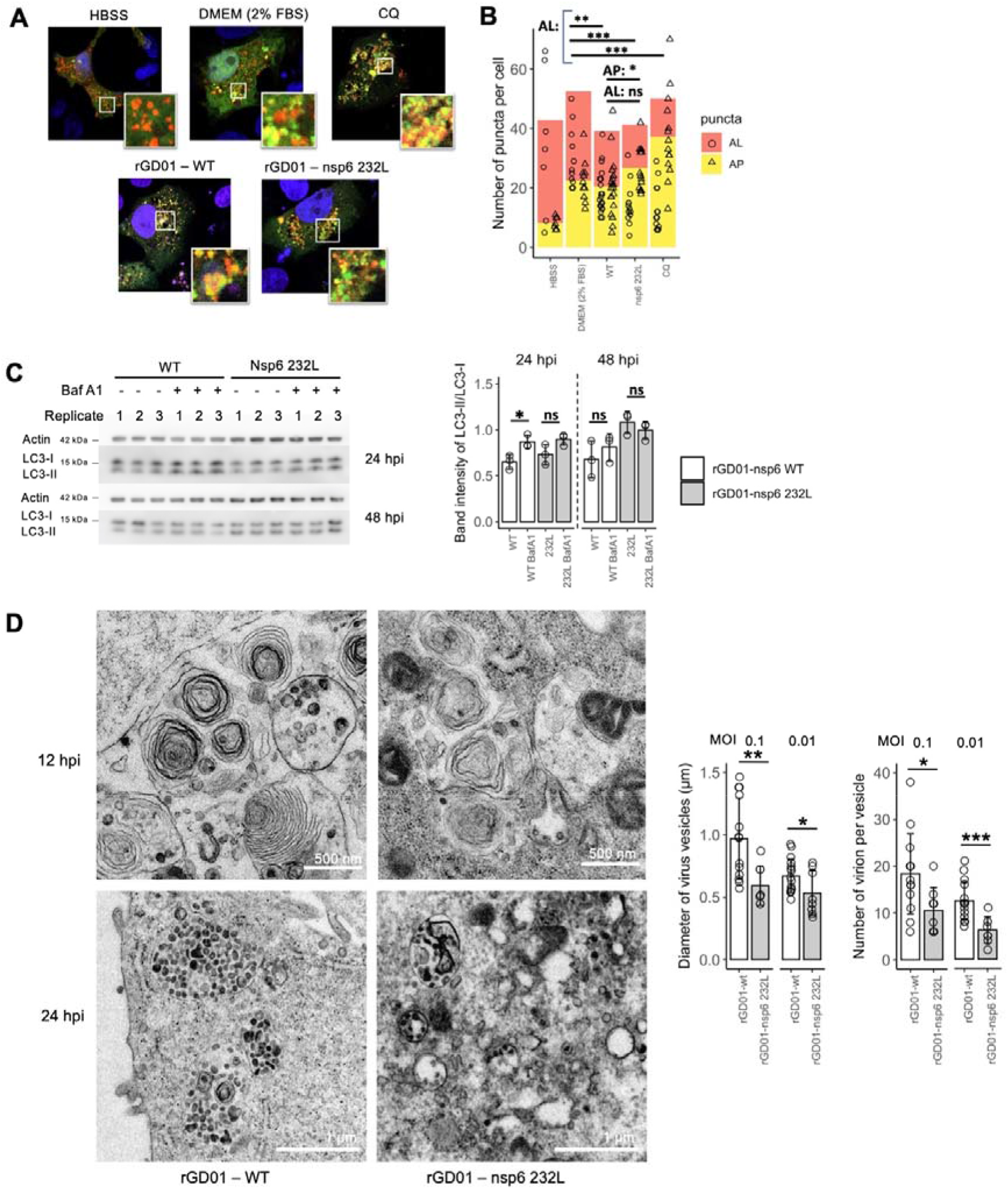
Nsp6 L232F mutation modulates autophagy. (A) Vero cells transfected with mcherry and EGFP tagged LC3 were infected with each of the recombinant viruses at MOI=0.01 and processed for immunofluorescence at 24 hpi. Hanks’ Balanced Salt Solution (HBSS) induced cell starvation and was used as a high autophagic flux control. DMEM supplemented with 2% fetal bovine serum act as the basal autophagic flux control. Chloroquine (CQ, 100 mM) inhibits AP fusion with AL as low autophagic flux control. Images are representative of at least three replicate experiments. (B) Summary statistics of the number of AP and AL puncta counted. Number of cells analyzed: HBSS (n=7); DMEM 2% FBS (n=11); WT (n=20); nsp6 232L (n=14). (C) Western blot of LC3-I/II in Vero cells infected at MOI=0.01. Bafilomycin A1 (BafA1, 0.1 μM) was added to cells for 2 h before harvest for protein lysate. Band intensity ratios of LC3-II/I were measured using ImageJ. Assays were performed in triplicates. (D) Electron microscopy of infected Vero cells at 12 hpi (MOI=1) and at 24 hpi (MOI=0.01). Diameters of virus vesicles and virion numbers per vesicle were measured from 24 hpi images using ImageJ. Number of cells analyzed: WT at MOI 0.1 (n=13), at MOI 0.01 (n=16); nsp6 232L at MOI 0.1 (n=8), at MOI 0.01 (n=8) Statistical tests were done using two tailed Student’s t test: p ≥0.05 (ns); p <0.05 (*); p <0.01 (**); p <0.001 (***).

### MERS-CoV nsp6 zippers endoplasmic reticulum, but L232F mutation did not affect zippering activity

A nsp6 ΔSGF 106-108 in SARS-CoV-2 was demonstrated to induce an increase in zippering of endoplasmic reticulum (ER) [21]. As CoV nsp6 shares a high degree of homology among coronaviruses, we hypothesised that the nsp6 L232F may affect the zippering activity in ER. nsp6 L232F is expected to position at the protein c-terminal cytoplasmic domain (Fig. 6A). Taking an experimental approach similar to that of Ricciardi et al. [21], we confirmed the ER zippering activity of MERS-CoV nsp6 from the punctate signals of ER reporter protein using a GFP fused with a cytochrome b5 C-terminal tail (GFP-cb5) (Fig. 6B). We further confirmed only N-terminal tagged MERS-CoV nsp6 will retain its native localization and zippering in ER (Fig. 6C). However, upon comparison of drug-induced expression between GFP-nsp6-wt and GFP-nsp6-F232L, no significant differences in the number of puncta structures per cell were observed at 1,3,6 and 8 hours post-induction, suggesting minimal impact of the nsp6 L232F mutation on ER zippering (Fig. 6D).

**Figure 6.**
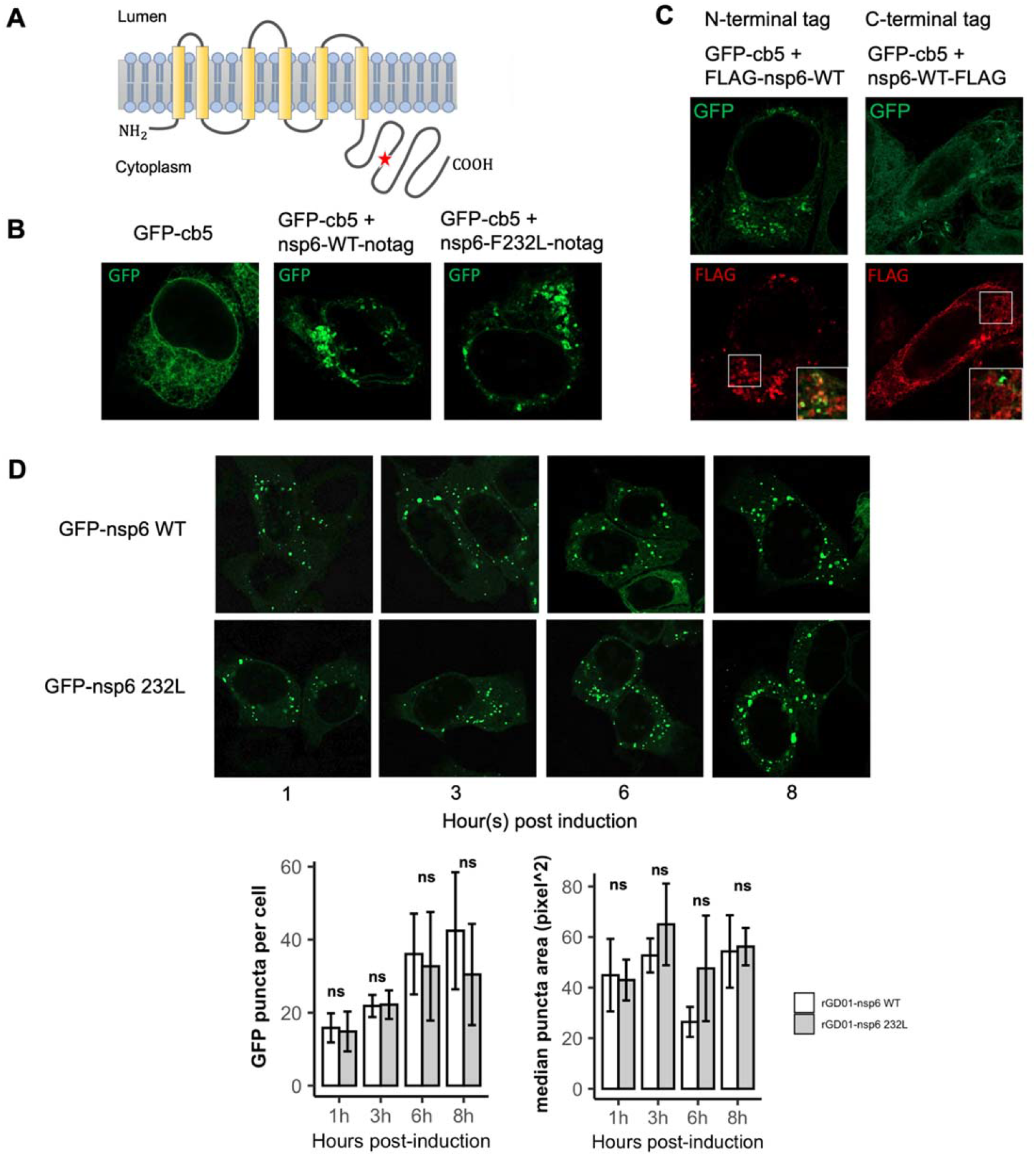
MERS-CoV nsp6 induces ER zippering but nsp6 L232F mutation did not affect ER zippering activity. (A) Illustration of the topology of MERS-CoV based on a previous study of murine coronavirus nsp6 from Baliji et al. [35]. The position of the L232F mutation is indicated as the star icon. (B) Hela cells expressing an ER reporter protein using a recombinant GFP with a cytochrome b5 hydrophobic sequence (GFP-cb5) or co-expressing with nsp6 WT or nsp6 232L without tag. GFP puncta indicate zippered ER structures. (C) Hela cells co-expressing GFP-cb5 with either a N-terminal or C-terminal FLAG tagged nsp6 WT. Insets, merged image of boxed areas. (D) Analysis of ER zippering formation in Hela cells expression GFP-nsp6 WT or GFP-nsp6 232L induced by doxycycline. GFP puncta per cell and area of puncta were analysed from images using ImageJ. Each setup was analysed with a cell number ranged from n=35-59. All images are representative of at least three samples. Statistical tests were done using two tailed Student’s t test: p ≥0.05 (ns).

## Discussion

Inter-species transmission of viruses may be associated with the emergence of adaptive mutations. There have been repeated zoonotic introductions of MERS-CoV from camels to humans. In this study, we aimed to identify adaptive mutations of MERS-CoV and understand their mechanistic basis. We identified nsp6 L232F mutation as one observed preferentially, almost exclusively, in humans. This mutation emerged independently in multiple lineages (lineages 2, 3, 4 and 5) in human MERS-CoV without being present in the phylogenetically related camel viruses, evidence of convergent evolution in humans. So far, only one camel MERS-CoV has been shown to carry this L232F mutation (Genbank accession number: KT368870.1), and this was a clade B virus. We used NGS analysis to demonstrate that nsp6 F232 was not present as a minor mutant population in camel nasal swabs. This suggested that the mutation likely appeared after inter-species transmission to humans although one cannot exclude the possibility that the mutation may be found rarely in camel viruses.

Replication of isogenic viruses pair (rGD01-wt / rGD01-nsp6-232L) demonstrated that rGD01-wt virus had significantly higher replication competence in-vitro and in ex-vivo cultures of human nasal and bronchial tissues. These data suggest the nsp6 L232F may confer a higher intrinsic replication efficiency in the human upper and conducting airways, though not in the lung. In hDPP4 knockin mice experimentally infected in-vivo with these isogenic viruses, rGD01-wt showed significantly higher viral titres in mouse lung. This suggest that the mutation is associated with an overall increase in virus production in vivo. Taken together, these findings may indicate that the nsp6 L232F mutation emerges in the human respiratory tract following zoonotic transmission as an adaptive mutation that confers replication advantage in the human respiratory tract. In the absence of a more relevant experimental model, we cannot estimate the impact of this mutation on transmission between humans.

In order to define molecular mechanisms underlying the nsp6 L232F mutation phenotype, we studied innate immune antagonism, autophagy and DMV formation. We found no significant difference in expression of a number of type-I interferon downstream genes following infection with the pair of isogenic viruses in Calu-3 cells. In the hDPP4 knockin mouse, where we observed the higher viral titer in the mouse lung from rGD01-wt infection, we found higher production of inflammatory cytokines such as IFNγ, MCP-1, and IP-10 (Supplementary Fig. 2). But this may merely reflect a consequence of the higher viral replication observed with the rGD01-wt virus rather than being a cause of this phenotype. Taken together, these data suggest the host innate immune responses were proportionate to the viral load and nsp6 L232F did not alter innate immune responses.

MERS-CoV has been shown to restrict autophagy during infection and ectopic expression of nsp6, Orf4b and Orf5 individually was shown to lead to autophagy restriction [23]. We demonstrated that nsp6 L232F mutation will reduce autophagic restriction in Vero cells during infection. Earlier studies have shown macro-autophagy is dispensable for MHV and SARS-CoV-1&2 replication [24-26]. This suggests a counteractive role of autophagy against viral replication. Since rGD01-WT replicated to higher viral titers than rGD01-nsp6 232L in Vero cells, lower autophagic restriction could be a host response for a more robust viral protein degradation though autophagy. Our EM images also revealed structures of multivesicular bodies upon infection. It has been shown that beta-coronaviruses can utilize lysosomes for egress instead of biosynthetic secretory pathway [27]. The larger virus vesicle size and higher virion numbers per vesicle supported a higher viral production of rGD01-nsp6 232L. It will be of interest to investigate whether nsp6 plays a role in the exocytic egress of MERS-CoV.

A recent study showed SARS-CoV-2 nsp6 with the ΔSGF 106-108 was shown to have a higher zippering activity than ancestral Wuhan strain nsp6, with a link to DMV formation [21]. We compared the zippering activity between GD01-nsp6-wt and GD01-nsp6-232L and identified no significant differences between them. The nsp6 L232F mutation is located in the c-terminal cytoplasmic tail of the nsp6, while the ΔSGF 106-108 in SARS-CoV-2 nsp6 is located in the hydrophobic transmembrane domain. The difference in the position of mutation may account for the difference in zippering activity results. Overall, the mechanistic role of how nsp6 mutation affects MERS-CoV replication requires further investigation.

There are some limitations to this study. First, we did not address the impact of the nsp6 L232F mutation on virus pathogenicity. Although there are mouse models that manifest lethality upon MERS-CoV infection, each have limitations and are not ideal to study the pathogenicity of a putative nsp6 L232F mutation effect. Introducing the nsp6 L232F mutation in mouse adapted strains which are pathogenic in mice is unlikely to be physiologically relevant. Second, we do not have an example where the nsp6 L232F mutation occurred within an individual. This would require serial sampling and sequencing of MERS-CoV viruses from very early in the zoonotic infection and such data is not available. Our findings provide a reason to initiate such systematic studies in zoonotic infections. The nsp6 L232F mutation was present in all viruses (n=34) associated with the outbreak of MERS-CoV in South Korea in 2015, including in the index case of that case-cluster [28-31]. It is worth noted that multiple generations of human MERS transmission did not result in reversal of F232 to L232 residue. The index case likely acquired virus infection through travel in the Arabian Peninsula and it is not clear if he was infected directly from a camel or if he acquired infection from another infected person. Another analysis of the MERS-CoV sequences from a hospital outbreak in Saudi Arabia (2018, n=18) showed the same leucine genotype at the nsp6 232 position [32]. This suggests that the nsp6 L232F mutation is not an indispensable human adaptive mutation. However, the phylogenetic evidence provides strong indirect evidence that this adaptation is occurring repeatedly in human infections. Finally, we did not examine the impact of the nsp6 L232F mutation on viral fitness in camels.

In summary, our study demonstrated that nsp6 L232F mutations are associated with interspecies transmission of MERS-CoV in humans and leads to higher replication competence in the human respiratory tract. These findings highlight the need for continuous surveillance in camels and humans to identify potential genotype changes relevant to human adaptation of MERS-CoV.

## Methods

### Cells

Vero cells (ATCC CCL-81) and Hela cells (ATCC CCL-2) were maintained in Dulbecco’s modified Eagle medium (DMEM), supplemented with 10% fetal bovine serum (FBS), 25mM HEPES and 1% penicillin with streptomycin at 37 °C, 5% CO^2^. Calu-3 cells (ATCC HTB-55) were maintained as above with the addition of 1% non-essential amino acid in the growth medium.

### Mutation analysis and phylogenetic analysis

MERS-CoV sequences (full or semi-full genomes >20 kb) were retrieved from Virus Variation, NCBI database. Sequences were first aligned by MAFFT and separated into either camel or human host for each individual ORFs. Mutation counting was implemented using custom script in R. Phylogenetic tree was generated using IQ-tree using its substitution model fitting [33].

### Reverse genetics and rescue of recombinant viruses

The bacterial artificial chromosome plasmid (pBAC) construct of infectious MERS-CoV/China01(GD01) strain (GenBank accession no. KT006149.2) was kindly provided by Prof J. Zhao (Guangzhou Medical School, China). MERS-CoV GD01 contains nsp6 L232F. An independent risk assessment of the genetic modification of MERS-CoV in this study was implemented in the Safety Office, the University of Hong Kong. The experiment envisaged is a loss-of-function mutation as we start from a human virus possessing the putative “human adaptive mutation” to mutate nsp232 to the camel genotype. To generate a recombinant GD01-nsp6 232L, a point mutation was introduced into the pBAC using a strategy of PCR mutagenesis and Gibson assembly. Primers were designed to amplify the whole genome of pBAC in long amplicon fragments of a size of 3 – 9 kb pairs each with 50 base pair overlapping regions. Specific forward and reverse primers containing the mutation were designed and used to amplify mutation carrying amplicons from the pBAC using PrimeSTAR Max DNA Polymerase (Takara). Each amplicon was gel-purified and assembled into a circular form using Gibson Assembly reaction mix (New England Biolabs) according to the manufacturer’s protocol. The ligated reaction mix was then transformed into MegaX DH10B T1R electrocompTM cells (Thermo Fisher) using electroporation. Transformed cells were then recovered and screened for positive clones by antibiotic selection. Positive clones were screened by a PCR reaction and sub-cultured for maxi-prep using NucleoBond Xtra Maxi EF kit (Macherey-Nagel) according to the kit instruction. Primers used are listed in the supplementary information.

Rescue of both wildtype and mutant virus were carried by transfection of pBAC infectious clones into Vero cells using lipofectamine 2000 (Thermo Fisher). Six hours post transfection, the transfection medium was removed and supplemented with 2% FBS supplemented DMEM. Success of virus rescue was confirmed by the observation of cytopathic effect 72 hours post-transfection. Rescued viruses were plaque purified in Vero cells and further sub-cultured to generate virus stocks, which were aliquoted and stored in -80°C until experimental use. Genetic identity of the virus stocks was confirmed by NGS. Titer of each stock of virus was determined by plaque assay.

### Replication kinetics in vitro

Calu-3 or Vero cells were seeded in 24-well tissue-culture plates at 1.5 × 10^5^ cells per well in 10% FBS supplemented DMEM. Prior to infection, cells were washed with PBS and serum-free medium was added to the cells. Infection of MERS-CoV was done at a multiplicity of infection (MOI) of 0.01 or 2 as indicated. After 1 hour of virus adsorption at 37°C, the virus inoculum was removed and cells washed with PBS twice to remove the remaining virus inoculum. Wells were re-filled with 2% FBS supplemented DMEM and incubated. Viral titres in the culture supernatants were quantified by the TCID_50_ assay.

### Growth competition assay

Vero cells were seeded with 1 × 10^6^ cells per well in a 6-well tissue-culture plate. Ratios of wildtype vs. mutant virus at 9:1, 1:1 and 1:9 were prepared and added to cells at MOI 0.01. After 72 hours of incubation at 37°C, virus medium was harvested for viral RNA extraction and aliquoted for subsequent passage. For subsequent viral passage, 10ul of virus medium was added to new plates seeded with cells supplemented in 2ml of 2% FBS supplemented DMEM and incubated another 72 hours of 37°C incubation. The procedures were repeated until passage 3. Viral RNA was extracted and sequenced across the mutation site by Sanger sequencing.

### Ex-vivo cultures of nasal, bronchia and lung tissues

Nasal, lung and bronchial tissues from healthy parts of lung removed as part of routine surgical care, excess to diagnostic requirements, was used for these studies. These experiments were approved by the Institutional Review Board of the University of Hong Kong/Hospital Authority Hong Kong West Cluster (UW-13-104 and UW-20-862). Tissues from at least three separate donors were separately infected with the wildtype or mutant virus by incubating the tissue fragments in 1 mL of each virus at a dose of 10^6^ TCID_50_/mL for 1 hour at 33°C for nasal tissues and at 37°C for bronchus and lung tissues. The tissue fragments were then washed three times with PBS and replenished with cell culture medium DMEM and incubated further for 72 hours. Culture supernatants were harvested at the timepoints as indicated and quantitated for quantification of live virus using a TCID_50_ assay. The methods have been previously described in detail [34].

### Experimental infection of human DPP4 knockin mice

6-8 weeks old human DPP4 knockin C57BL/6 mice (kindly provided by Prof. Stanley Perlman, University of Iowa) were anesthetized and infected intranasally with 10e^4^ pfu of wildtype or mutant virus in 25 μl of PBS. Mock control mice were similarly treated with PBS only. Four mice from each virus group were euthanised at 4 hours, day 1, day 2, day 3 and day 5 post infection. The mouse lungs were removed and homogenized in 1 ml PBS solution and stored at - 80 °C until use. Supernatants of centrifuged homogenate were tested for viral titers for TCID_50_ assay in Vero cells.

### qPCR of innate immune gene expression

Virus infected cells were washed gently with 1ml of PBS twice. Cellular RNA was extracted and reverse transcribed into cDNA using random hexamers and measured for gene expression using qPCR assay. Primers used are listed in the supplementary information.

### Autophagic flux assay and LC3 immunoblotting

The effect of MERS-CoV infection on autophagic flux was determined by the transfection of a pHR’-CMV-eGFP-mCherry-LC3 plasmid (provided by Dr. Sumana Sanyal, University of Oxford) encoding fluorescent LC3 which emits as yellow puncta in autophagosomes and as red puncta in autolysosomes. Chloroquine (Sigma-Aldrich, C6628) and Hanks’ Balanced Salt Solution (HBSS) (Thermo Fisher) were used as reduced and increased autophagic flux controls respectively. Vero cells were transfected with pHR’-CMV-eGFP-mCherry-LC3 using Lipofectamin 2000, followed by MERS-CoV infection at MOI 0.01. Cells were fixed 24h post infection by 4% PFA and processed for confocal imaging.

Cells were extracted for protein using RIPA buffer with freshly supplemented protease inhibitors. Protein samples were subsequently added with sample buffer and incubated at 95°C for 10 mins. Western blotting of LC3-B was labelled with anti-LC3-B primary antibodies (1:1000, Cell Signaling Technology, #3868) and was normalized with anti-Actin antibodies (1:4000, Thermo, MA1-744). Band intensities of LC3B-I and LC3B-II were measured in Fiji software (ImageJ, National Institutes of Health) and normalized to band intensity of Actin.

### Electron microscopy

MERS-CoV infected Vero cells were harvested at 9 hpi (MOI=1) and 24 hpi (MOI=0.01). Cells were washed once with PBS and scrapped from well-plate in 10% formalin. Cells were centrifuged down into a pellet and fixed overnight. In the next day, cells were fixed in 2.5% glutaraldehyde in cacodylate buffer (0.1 M sodium cacodylate-HCl buffer pH 7.4) overnight, followed by processing, sectioning and staining for transmission electron microscopy using Philips CM100.

### Plasmid construct for immunofluorescence study

Nsp6 gene fragments (WT and nsp6 232L) were amplified from the pair of recombinant pBAC-GD01 and cloned into a pCAGGS vector containing a FLAG tag either at the N-terminal or C-terminal, or without any tag using Gibson Assembly system. Inducible vectors were generated by cloning the nsp6 gene fragment into a doxycycline-inducible pCW vector containing a GFP tag at the N-terminal. Empty pCW was kindly provided by Alessia Ciarrocchi & Gloria Manzotti (Addgene plasmid # 184708). ER reporter protein GFP-cb5 was constructed by amplifying GFP with oligos containing the cytochrome b5 transmembrane domain sequence (5’-ITTVESNSSWWTNWVIPAISALVVALMYRLYMAED-3’) and cloned back into the pCAGGS vector using Gibson Assembly system.

### Nsp6 expression in ER and immunofluorescence staining

Hela cells were seeded on a cover glass placed in a 12 well plate. In the next day, cells were transfected with pCAGGS-nsp6, pCAGGS-FLAG-nsp6, pCAGGS-nsp6-FLAG, or pCAGGS-GFP-cb5 using TransIT-LT1 (Mirus Bio) according to the manufacturer’s protocol. At 24h post transfection, cells were fixed in 4% paraformaldehyde and processed for immunofluorescence staining. Cells were permeabilized with 0.1% Triton X-100 for 10 minutes and blocked with normal goat serum for 1 h at room temperature. Nsp6 expression was stained using a primary rabbit polyclonal anti-FLAG (1:100, Bethyl, A190-102A) and a secondary goat anti-rabbit IgG H&L (1:200, Alexa Fluor® 647), visualized in confocal microscopy (LSM980, Zeiss).

### ER zippering activity

Hela cells seeded on a cover glass were transfected with pCW-GFP-nsp6 or pCW-GFP-nsp6-232L using TransIT-LT1 according to the manufacturer’s protocol. At 24h post transfection, expression of nsp6 was induced by adding 1 μg ml^−1^ doxycycline (Sigma-Aldrich). Cells were fixed at 1h, 3h, 6h and 8h post induction by 4% paraformaldehyde and analyzed by confocal microscopy.

### Confocal microscopy and image analyses

Cells were imaged on a Zeiss LSM980 system. Images from the experiment were taken under the same laser parameter settings and magnification. Puncta from the images were counted and measured using Fiji software (ImageJ, National Institutes of Health) under the same colour threshold settings across images.

### Statistical analysis

Comparisons were performed using two-sided unpaired Student’s t-test. All the data using ex-vivo tissue cultures were compared using paired t-test. P values of <0.05 were considered statistically significant.

## Data availability

All genomic sequence data are publicly available data through NCBI Genbank (https://www.ncbi.nlm.nih.gov/genbank/). Accession numbers of viral sequences used in the phylogenetic analysis are listed in the supplementary information. Analysis scripts for mutation analysis and sequencing data can be accessed at Github depository. Source data are provided with this paper. Correspondence and requests for materials should be addressed to Malik Peiris, E mail

## Acknowledgements

We acknowledge Prof Stanley Perlman for providing critical comments on results of this study and Dr. Eric H.Y. Lau for advice on statistical analysis. We acknowledge Mr. Garrick Yip and Dr. Tomas Lyu for the technical discussion on generating the mutagenesis in pBAC. We acknowledge the Centre for PanorOmic Sciences, the University of Hong Kong, for the technical support of the confocal system. The research was funded from a research grants from the Health and Medical Research Fund of the Government of the Hong Kong Special Administrative region (No 19181032) and from the US National Institutes of Health (contract no. U01-Grant AI151810).

## Author information

### Contributions

RTYS and MP conceived and planned the overall study while LLMP and SS contributed to design of virology and cell biology experiments. RTYS and DKWC carried out the phylogenetic analysis. KPYH, JCWH, MCC, KCN, HWY & MCWC carried out the ex-vivo cultures of human respiratory tissues. CKPM carried out mouse infection experiments. JMN carried out and interpreted electron microscopy studies. RTYS wrote the original draft. RTYS, MP and SS reviewed and edited the manuscript.

### Corresponding author

Malik Peiris

